# Development and field test of an intervention to reduce conflict in faculty-doctoral student mentoring relationships

**DOI:** 10.64898/2026.01.29.702507

**Authors:** Trevor T. Tuma, Emily Q. Rosenzweig, Justin A. Lavner, Yichi Zhang, Erin L. Dolan

**Affiliations:** Department of Biochemistry and Molecular Biology, University of Georgia, Athens, GA 30602; Department of Human Development, Teachers College, Columbia University, New York, NY 10027; Department of Psychology, University of Georgia, Athens, GA 30602; Department of Educational Psychology, University of Georgia, Athens, GA 30602

**Keywords:** Mentorship, Graduate education, Professional development, Conflict, Attributions

## Abstract

Mentoring is a critical component of graduate education. However, conflicts can occur between faculty mentors and their graduate students, which can undermine the quality of these relationships. We leveraged attribution theory and relationship science to develop a novel professional development intervention that combines attribution retraining to enhance faculty beliefs that they can improve their mentoring relationships, and conflict management training to build faculty skills in having productive problem-solving conversations with their graduate students. We piloted and refined the intervention, then conducted a field test of the intervention with life science faculty (n = 71) from U.S. universities. Participants were randomly assigned to an asynchronous self-guided condition or to a self-guided + synchronous facilitated peer discussion condition. We measured faculty beliefs, perceived skills, and self-reported behaviors when encountering conflicts before and after participating in the intervention. Faculty in both conditions reported significant reductions in the frequency of conflicts with their students, the time and energy they spent addressing conflicts, and the extent to which conflicts disrupted their research productivity. Faculty also expressed increased confidence that they could manage conflicts. Our results suggest that the intervention has the potential to improve faculty capacity to effectively navigate conflicts with their graduate students.

**Highlight summary:** A mentoring intervention for faculty combining attribution-retraining and conflict management skill-building strengthened faculty self-efficacy and motivational beliefs and reduced mentoring conflicts.

## INTRODUCTION

A life science PhD student’s relationship with their research advisor is critical to the quality and success of their graduate training experience (Sverdlik et al., 2018; Zhao et al., 2007). Research advisors are positioned to mentor their students by providing career and psychosocial support to help their mentees advance toward their professional and personal goals (Tuma & Dolan, 2024). Regretfully, mentoring relationships between PhD students and their advisors are not always positive (Tuma et al., 2021, 2025). Negative interactions with advisors undermine students’ professional development, career intentions, and well-being (Byrom et al., 2022; Sowell et al., 2015; Tuma et al., 2021). Accordingly, a major recommendation of a National Academies consensus committee on mentoring (2019) was to improve the mentoring experienced by graduate students.

Efforts to improve mentoring have produced a variety of mentoring workshops, programs, curricula, and how-to guides. A recent review highlighted >100 published reports of mentorship interventions (Gangrade et al., 2024), but noted that less than half were designed for faculty and most presented limited evidence of effectiveness. Furthermore, only one explicated a theory that informed the design of the mentoring intervention (Lewis et al., 2016, 2017). The absence of theoretical grounding for mentoring professional development introduces two limitations of this work. First, it can limit the potential for mentoring interventions to be effective if they do not draw sufficiently from prevailing ideas of how people learn or how behavior changes. Second, when studies of mentoring interventions lack theoretical grounding, they miss an opportunity to test ideas about why these interventions may (or may not) be effective. Based on this review, Gangrade and colleagues (2024) called for the development and rigorous testing of additional mentoring interventions.

The most widely disseminated mentoring professional development curriculum for research advisors, *Entering Mentoring*, focuses on fostering effective mentoring relationships by helping mentors learn about and practice six main competencies: aligning expectations, communicating effectively, addressing diversity, assessing understanding, developing independence, and promoting professional development (Pfund et al., 2015). Participation in this curriculum can improve both mentor and mentee perceptions of mentor competence in the target areas and behaviors (Pfund et al., 2014). In our experience implementing mentoring professional development, research advisors describe *Entering Mentoring* a useful starting point for learning how to mentor. Yet faculty express a desire for additional guidance on navigating conflicts that arise in their mentoring relationships with graduate students, which the curriculum does not explicitly address. These observations align with national reports highlighting variability in mentor preparation and a dearth of professional development on how to address complex relational challenges when mentoring students (National Academies of Sciences, Engineering, and Medicine [NASEM], 2019).

Graduate student-research advisor conflict is an inevitable and normal part of research mentoring relationships given the interdependence of graduate students and their research advisors and the academic, cultural, and personal diversity within STEM graduate education (Klomparens et al., 2008). How conflict is managed determines its effects (DeChurch et al., 2013) and has critical implications for graduate student success (Klomparens et al., 2008). If managed well, conflict can lead to improvements in goal clarity, creativity, personal growth, and trust in the relationship (Klomparens et al., 2008; Wheelan, 1994). Unfortunately, most research advisors are not formally trained to appraise, manage, and resolve conflicts productively, increasing the likelihood that conflict will be managed poorly (Lee et al., 2015; NASEM, 2019). This can lead to negative consequences for graduate students, including loss of confidence, loss of funding, greater time to degree, and attrition from their graduate program or the field altogether (Brockman et al., 2010). Yet existing interventions provide little explicit guidance to support research advisors in productively navigating these conflicts. In fact, Gangrade and colleagues (2024) noted that learning to navigate conflicts was largely absent from published reports of mentoring interventions.

Helping research advisors address conflicts with their graduate students is further complicated by advisors’ endorsement of counterproductive beliefs about mentoring and students. For example, faculty may believe that effective mentoring is about finding the right match (Tuma et al., 2021; Tuma & Dolan, 2024); that success in STEM fields requires brilliance, natural talent, or intellectual giftedness (Hernandez et al., 2025; Leslie et al., 2015); or that mentors should prioritize controlling who continues in research over providing support (Tuma et al., 2021). These types of beliefs may make research advisors less willing to invest time and effort in mentoring professional development, because they may not think they are contributing to conflicts with mentees or that their ability to become better mentors can be improved (Weiner, 1985).

Collectively, this research suggests that interventions are needed to support research advisors in developing the skills to navigate conflicts effectively. Furthermore, such interventions may be more effective if they are coupled with activities that help research advisors believe they can successfully address conflicts through their own actions. In this study, we present the development and field testing of a professional development intervention for faculty research advisors^1^ designed to incorporate these two ideas. The intervention, Improving Mentorship Practice through Attributions and Conflict Training (IMPACT), combines activities to help mentors feel more capable of addressing conflicts with PhD students (i.e., a motivational intervention technique called attribution retraining) and activities to teach mentors skills to have more productive conversations with their students about these conflicts (i.e., conflict management skills training). The attribution retraining component draws from theory and best practices related to motivational beliefs, and the conflict management conversations component draws from principles and best practices from relationship science.

## THEORETICAL AND CONCEPTUAL BACKGROUND

The IMPACT intervention has two main, theoretically informed components: (1) helping mentors feel capable of addressing conflicts with students, and (2) teaching mentors the skills to have better conversations to address conflicts with students. Next we describe the theory, principles, and practices for each component and how we applied this work in the design of IMPACT.

### Helping Mentors Feel Capable of Addressing Conflicts

The first major component of IMPACT aimed to motivate mentors to invest time and effort to learn conflict management skills. For individuals to learn and use conflict management skills, they must first believe that they are capable of addressing conflicts with those skills. That is, they need to have *self-efficacy,* defined as the perception of being capable of engaging in a designated course of action (Bandura, 1997). Individuals with higher self-efficacy about providing instruction, mentorship, or leadership are more successful in doing so (Astrove & Kraimer, 2022; Dwyer, 2019; Lazarides & Warner, 2020).

The attribution theory of motivation (Graham, 2020; Weiner, 1985) offers an explanation of how mentor self-efficacy for addressing conflicts can develop. This theory delineates how an individual’s beliefs about the causes of past events strongly shape their self-efficacy for engaging in future activities. Attribution theory posits that individuals can make three types of attributions about the cause of a past event: (1) Locus, defined as whether the event was caused by oneself or something external (internal vs. external); (2) Controllability, defined as whether the individual causing the event was in control of their behavior (controllable vs. uncontrollable); and (3) Stability, defined as whether the same cause-effect relationship would happen again (fixed vs. malleable). For instance, if students make fixed, external attributions about an academic failure (e.g., they think they failed an exam because of an instructor’s teaching style as opposed to their own choice of study strategies), they will report lower self-efficacy for addressing academic challenges in the future (Graham & Taylor, 2022). This is because they believe that changing their own behavior is not likely to change anything moving forward, given that the situation was not caused by their own actions. In contrast, if students attribute an academic failure to controllable causes that are malleable across situations (e.g., how and when they choose to study), they are likely to perceive higher self-efficacy for the next attempt because they believe changing their behavior can lead to better outcomes (Graham, 2020).

When applied to mentoring, attribution theory suggests that mentors who view conflicts with students as controllable and malleable are more likely to report greater self-efficacy in their ability to address them. For example, if a mentor has a conflict with a student and decides it is due to something unchanging about the student (e.g., they think the student is hypersensitive to criticism), the mentor may perceive that an intervention to develop their own skills is not going to help address that issue. Thus, the mentor would be less likely to invest the time and effort to learn and improve their conflict management skills. Conversely, if a mentor believes that the conflict was due, at least in part, to something they could control (e.g., how they talked with the student about the conflict), then they will be more likely to invest the time and effort to learn new skills to address conflicts in the future because they will be more confident the skills will help. Importantly, we do not mean to imply that mentors should blame themselves (or their students) for conflicts—the reality of most mentoring conflicts is that they are multi-faceted, involve contributions from both parties, and arise from complex causes. However, attribution research suggests that encouraging mentors to recognize their own abilities to address conflict situations, even when other causes are also at play, could help them feel more capable of addressing conflicts and ultimately be more successful at doing so.

The IMPACT intervention used techniques from attribution retraining interventions to help mentors attribute the causes of conflicts as internal, controllable, and malleable, with the aim of fostering their self-efficacy for addressing conflicts with their students. More than 30 years of research has shown that attribution retraining interventions help students attribute their academic challenges to malleable and controllable causes as a path to promote their self-efficacy for learning in the face of challenges (see Graham & Taylor, 2022; Siegel Robertson, 2000, for reviews). Growth mindset interventions are also grounded in attribution theory, helping individuals attribute their intellectual ability to things they can control (see Yeager et al., 2019 for review). Participants in both types of interventions typically complete tasks that emphasize the importance of effort or the use of improvement strategies to remind them that they are capable of addressing academic challenges. For example, an intervention for students might have participants read others’ testimonials about how they learned that their challenges in a course were due to insufficient effort as opposed to low ability. Then they might write a reflective essay about one of their own academic failures that was due to low effort. These types of interventions have been highly successful at changing students’ attributions, improving grades, enhancing classroom well-being, and boosting effort in school (Blackwell et al., 2007; Hamm et al., 2020; Perry et al., 2010; Yeager et al., 2019). To date, prior work has also used attribution retraining and growth mindset intervention techniques to improve qualities of other dyadic relationships, including teacher-student, employer-employee, and peer-to-peer relationships (Bryan et al., 2021; Heslin et al., 2005, 2006; Hudley et al., 1998). A recent study applied attribution retraining techniques in mentoring relationships, but the intervention was aimed at improving student mentees’ beliefs (Tuma et al., 2025). Despite the power and potential of attribution retraining interventions, we know of no prior studies that have leveraged attribution retraining to help research advisors improve as mentors.

Attribution retraining adds additional potential value to mentoring professional development because it is theorized to tap into self-sustaining belief-behavior cycles. These cycles, known as recursive psychological processes, can promote long-term changes by continuing to affect an individual long after the intervention ends (Walton, 2014; Walton & Wilson, 2018). For example, an intervention can change a belief, such as self-efficacy, which in turn promotes an adaptive behavior, such as effort, which provides reinforcement that further changes the belief (i.e., greater self-efficacy as a result of investment of effort), leading to even more adaptive behavior (i.e., more effort), and so on. We used attribution retraining in the IMPACT intervention to promote enduring changes in mentors’ beliefs and behaviors.

### Skills to Promote Productive Conversations about Conflicts

Changing mentors’ attributions about conflicts with their graduate students is a critical but insufficient step to improve mentoring relationships. Mentors must also be equipped with useful skills they can deploy effectively when conflicts arise. Thus, the second major component of the IMPACT intervention aimed to teach mentors skills to have more productive conversations to address conflicts with their graduate students. Research shows that effective conflict management skills can positively impact relationship quality in a variety of interpersonal relationships, including employee-employer (e.g., Gilin Oore et al., 2015), parent-child (e.g., Openshaw et al., 1992), and friendship (e.g., Chow et al., 2013) relationships. A host of conflict management training interventions have been developed to improve these skills, targeting a range of relational contexts such as healthcare professionals (e.g., Berkhof et al., 2011), college students (e.g., Davidson & Versluys, 1999), and romantic couples (e.g., Worthington Jr et al., 2015). Among couples, for example, behavioral interventions focus on teaching partners how to share their thoughts and feelings effectively, engage in active listening, and use a stepwise process to address conflicts and other challenges (Epstein & Baucom, 2002). These steps include identifying the issue, clarifying why the issue is important and what each partner’s needs are, discussing possible solutions, deciding on a solution, and testing out the solution during a trial period. Very little research has evaluated conflict management skills training with research advisors and graduate students, though the limited work that has been done with graduate students (Brockman et al., 2011; Schaller & Gatesman-Ammer, 2022) and medical residents and academic health care faculty (Zweibel et al., 2008) suggests some benefits.

The IMPACT intervention used techniques from this prior work to teach mentors a three-step process for addressing conflicts with their graduate students: (1) Discover, (2) Discuss, and (3) Decide (Epstein & Baucom, 2002). The Discover step aimed to help mentors discover their own and their student’s point of view about a problem to better understand what is at the core of a conflict and why it matters to them personally. In the Discuss step, mentors learned how to share their perspective on the problem with their student and listen to their student’s perspective in a way that allows the conversation to continue in a productive manner. In the Decide step, mentors learned how to work with their students to identify an initial solution to the problem and how to follow up to evaluate how the solution is working and revise if needed. The intervention purposely focused on the “process” (i.e., the conversation about problems) as well as the “product” (i.e., the solution to the problem) to equip mentors with an approach that could be applied to a broad range of conflicts that might arise in their mentoring relationships.

Strategies within each of these steps were informed by specific behaviors and general principles from previous work (e.g., Epstein & Baucom, 2002). For example, in the Discover step, mentors were led through a short reflective process to identify one single issue that is most concerning to them before approaching the student, consistent with guidance that complex problems should be broken into smaller ones and addressed one at a time (e.g., Epstein & Baucom, 2002). The Discuss step included strategies to help the mentor communicate clearly about the issue and listen effectively to their student. For example, mentors were encouraged to keep their remarks brief when they share their perspective with the student, to state their views subjectively, and to avoid global judgments (e.g., “you are always late”); these and other strategies keep the conversation focused on the problem at hand and make it easier for the student to respond. Active listening strategies were also reviewed, including paraphrasing to check for understanding, acknowledging the other’s thoughts and feelings through verbal and nonverbal messages, and asking questions with the goal of learning about the other’s experience (Stone et al., 2023). Active listening is an effective tool to mitigate, prevent, and resolve conflicts because it helps uncover the information needed for conflict resolution while also addressing a fundamental need for others to feel understood (Van Slyke, 1999). The IMPACT intervention also discussed strategies to manage challenges that may arise during these conversations, such as “taking a pause” when either the mentor’s or the student’s emotions are too intense or if the conversation has stalled. This pause allows both people to collect their thoughts and engage more productively when the discussion resumes (Mischel et al., 2014). Finally, the Decide step outlined strategies for the mentor to decide together with their student on an initial solution to the problem, such as identifying criteria for a solution (i.e., the rules the solution must meet), generating a list of possible solutions, and choosing a solution to try, as well as following up afterward to evaluate how things are going (Epstein & Baucom, 2002). Collectively, the IMPACT intervention aimed to foster communication that is clear, responsive, and task focused.

### The Current Study

In the present paper, we describe our process for developing and refining the IMPACT intervention as an example of a theory- and principle-based intervention. We also describe the results of a field test of IMPACT with 71 life science faculty as an initial assessment of the effectiveness of IMPACT. Specifically, we aimed to (1) assess the feasibility of implementing an asynchronous self-guided version of IMPACT and a combined asynchronous self-guided + synchronous peer discussion version of IMPACT and (2) determine whether participation in IMPACT influenced mentors’ motivational beliefs related to conflicts with their graduate students (i.e., self-efficacy and attributions) and their self-reported behaviors and experiences with such conflicts.

## METHODS

The present study was determined to be exempt by the University of Georgia’s Institutional Review Board (PROJECT00007308).

### Intervention Design

We used a design-based research process to develop IMPACT as a novel attribution-retraining and conflict management skill-building intervention for faculty mentors of life science PhD students. The intervention comprises 10 online modules that participants complete on their own over six weeks and a companion series of facilitated small-group discussions for peer discussion participants (i.e., six weekly synchronous sessions on Zoom).

We began by drafting the ten modules, starting with the modules that would support mentors in retraining their attributions. We sought to shift mentors’ thinking that their relationships with graduate students and conflicts with their students were controllable and malleable rather than uncontrollable or unresolvable. We drew from attribution retraining and belief reappraisal interventions to create text and activities for these modules (Hamm et al., 2020; Rosenzweig et al., 2020, 2022; Tuma et al., 2025; Walton & Cohen, 2007, 2011). Then we drafted the modules intended to help mentors learn the three-step process of Discover-Discuss-Decide for engaging in productive problem-solving conversations with their students. We developed these materials by drawing from the principles for effective problem-solving conversations described earlier, including identifying the issue and clarifying why it matters to the mentor, seeking to understand the issue from the student’s perspective, discussing possible solutions, deciding on a solution and testing it out, following up to gauge how the solution is working, and making adjustments as needed. We also integrated strategies for managing the emotional elements of navigating conflicts, including how to proceed when the student’s or mentor’s emotions get intense and how to interact after the conflict. Finally, we used findings from qualitative research about negative mentoring experienced by graduate students (Tuma et al., 2021) to inform the examples included in the modules.

We designed all module materials to align with evidence-based recommendations for promoting deep and generative learning (Fiorella & Mayer, 2016). We also sought to optimize engagement by designing IMPACT to be intensive enough to promote lasting skill acquisition but not so demanding that participants disengage due to perceived burden (Hawkins et al., 2008; Wade et al., 2014). Achieving this balance is essential for promoting long-term behavioral changes (Hamm et al., 2020; Segrin & Givertz, 2003; Walton & Wilson, 2018). Across all of the modules, we provided mentors with opportunities to engage autonomously by making choices about what to discuss and to think actively and deeply about the content. As one example, for some of the modules, mentors read quotations describing other mentors’ experiences dealing with conflicts with graduate students. To promote engagement and autonomous reflection in this task, we asked mentors to rate each quotation for its level of informativeness and how relevant it was to their experiences as a faculty member. We also prompted mentors to rank the quotations from most to least favorites and write a brief explanation about why their top-ranked quote was their favorite. Additionally, in the modules focused on conflict management skills, faculty were presented with an example scenario that continued through all of the modules so that they could see the skills in action and altogether. We integrated tasks throughout the modules for mentors to practice applying skills in their own mentoring interactions and write advice to a colleague using the principles in the modules. We designed these tasks to prompt further engagement and to leverage the notion of “saying-is-believing,” with the goal that mentors would endorse the messages in the intervention more strongly (Higgins & Rholes, 1978).

We developed two versions of IMPACT: (1) the self-guided version described above, which has the potential to be flexible and scalable because it requires no facilitation, minimal software infrastructure, and modest personnel effort to implement, and (2) a self-guided + facilitated peer discussion version, which requires greater coordination and facilitation but also offers additional opportunities for practice, feedback, and peer learning. To create the peer discussion version of IMPACT, we developed a set of materials to mirror the self-guided materials while providing additional opportunities for skills-based practice and feedback in small peer groups. We drew from Social Learning Theory (Bandura & Walters, 1977) and Communities of Practice Theory (Lave & Wenger, 1991) to inform the design of tasks for the discussions. Social Learning Theory posits that individuals develop new skills more effectively when they can observe others, try out behaviors, and receive feedback in real time. Communities of Practice Theory posits that individuals develop competence and confidence through participation in a community that shares a common task. Consistent with both theories, we designed the peer discussion materials for mentors to observe peers, articulate reasoning, try out new strategies for addressing conflict, and receive social support and guidance from the intervention developers. We included new scenarios on the same topics as in the self-guided materials, with prompts for reflection and discussion as well as role-playing activities, opportunities to ask questions, and other interactive exercises. We created a slide deck and detailed instructions for facilitators for each of the six peer discussions, along with any handouts. All peer discussions followed a similar structure: introduction and warm-up activity; articulation of learning objectives; brief review of the self-guided content; structured discussion and applied activities related to the week’s topics; and concluding highlights of key points for the week.

### Intervention Piloting and Refinement

We conducted iterative pilot testing to refine the self-guided modules after creating the initial materials and prior to field testing (Figure 1). We first collected feedback from three experts. The first expert was a faculty member in psychology with expertise in attribution retraining. This expert was asked to review a subset of the modules that addressed motivationally relevant content to improve alignment with attribution theory. They provided feedback during a Zoom meeting with a subset of the research team. The second and third experts were life science faculty members experienced in mentoring graduate students and leading graduate education programs. These individuals reviewed all of the modules and gave written and oral feedback through a series of asynchronous work sessions and Zoom meetings with a subset of the research team. They were asked to identify aspects of the content they found most and least compelling and to flag any language or scenarios they considered inauthentic or inapplicable to life science graduate education. Faculty were compensated $75 for each hour they spent reviewing the modules and completing the study activities. All sessions were audio-recorded, and two researchers wrote notes during each session. The entire research team then reviewed all feedback and revised the modules accordingly.

**Figure 1.**
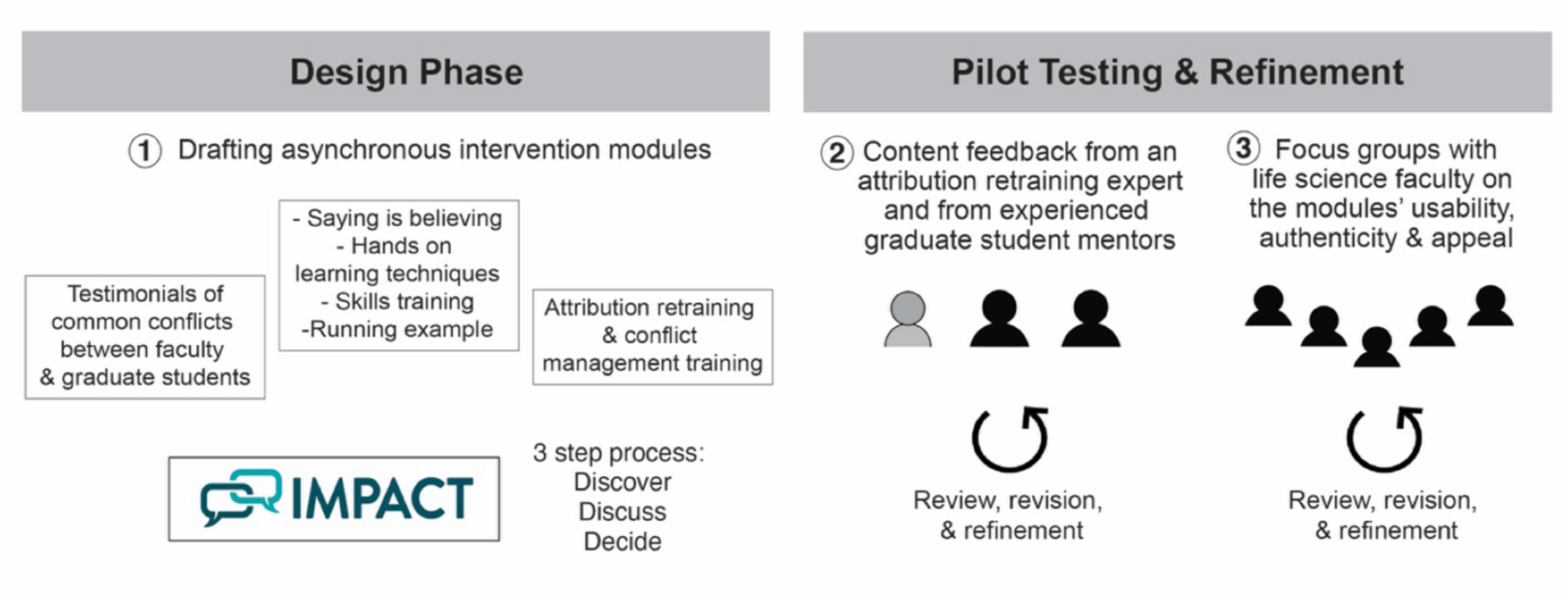
The design-based research process used to develop the IMPACT intervention. We adapted content, exercises, and tasks from previous attribution retraining and conflict management skill-building interventions. We refined the materials to be cohesive, relevant, and impactful for mentors of life science PhD students, including realistic mentoring conflicts characteristic of life science graduate student-faculty relationships as well as multiple opportunities for engagement and practice. After initial development, the content was reviewed by a subject matter expert in attribution retraining and by faculty with extensive experience mentoring life science PhD students. Their feedback was used to iteratively revise and refine the self-guided materials to align with the intended psychological principles while ensuring feasibility, authenticity, and appeal to life science faculty.

We then sought feedback through focus group discussions with life science faculty. We emailed the study information to faculty who were mentoring life science PhD students in research-intensive universities across the US. We strategically selected five faculty members from different universities who ranged in their rank and mentoring experience with at least three years as a faculty member and had experience advising at least two PhD students for at least one year each. The majority of participants identified as women (women = 4, men = 1), white (white = 4, prefer not to respond = 1), non-Hispanic (non-Hispanic = 4, Hispanic or Latine = 1), and continuing generation college graduates (continuing generation = 4, first generation = 1). Most participants were tenured (Assistant Professor = 1, Associate Professor = 4), had been faculty members for an average of ∼10 years (SD = 3), and had mentored graduate students for an average of 9 years (SD = 3). All participants had previously completed some mentoring professional development, although the scope and intensity of that training varied.

We asked participants to complete the modules on their own and then to provide feedback on content, clarity, and relevance of the module content during a series of Zoom meetings. Through this process, we sought to gauge how well IMPACT content aligned with the faculty members’ own mentoring experiences, what they thought they learned, what the reasoning was behind their responses, and whether they had any suggestions for improvements, especially in response to any language, concepts, or examples they found confusing, unrealistic, or discomforting. Again, the research team reviewed all feedback and revised the modules accordingly.

Several themes emerged from pilot testing (Table 1). First, participants noted that the focus on conflict in the early modules could be interpreted negatively and had the potential to be off-putting to participants who viewed themselves as effective mentors. Accordingly, we revised those modules to highlight the rewards and benefits faculty can realize from mentoring. We also revised the content to emphasize that conflicts are normal, with the goal of reassuring participants that encountering conflicts does not indicate poor performance or an inability to improve. Second, participants raised concerns regarding the authenticity of some of the conflicts depicted in the modules. To address this, we revised the scenarios to be more authentic and believable in life science research. We also updated terminology (e.g., replacing the term “mentor” with “PI”) to better reflect everyday language in life science research settings. Third, participants were eager to learn specific strategies for addressing conflicts earlier in the intervention. We consolidated and reorganized the material such that Week 1 foreshadowed the skills to be learned, and Week 2 began skills training, instead of starting training in a later week as we had originally planned. Finally, participants noted stylistic issues, such as advice being too prescriptive, examples being too simple, and content being too wordy. To address these concerns, we replaced numbered steps with more flexible guidance and illustrative examples. We also revised the quotations and example scenarios to be more complex and nuanced with less obvious resolutions, and we streamlined the text. The result of this process was a refined set of modules to be used in field testing of the IMPACT intervention.

**Table 1.** Changes made to module materials during iterative pilot work.

| Issues raised | Changes made to modules |
| --- | --- |
| **Negative tone:** Early focus on conflict seemed negative, potentially putting off faculty who see themselves as good mentors | • Changed introduction to focus on rewards of mentoring<br>• Repeatedly emphasized that conflict is a normal part of mentoring and that having conflicts does not imply anything “bad” is going on |
| **Lack of authenticity:** Examples did not always highlight or attend to life sciences-specific concerns (e.g., lab safety conflicts not being open to debate) | • Specifically vetted all examples in focus groups for suggestions to make the examples precise and specific to life sciences. Added qualifiers where needed to address issues that were raised (e.g., “this did not present a safety issue, so...”)<br>• Changed term “mentor” to “PI” to emphasize life sciences terminology |
| **Need to further emphasize the importance of addressing conflicts:** Examples could emphasize why avoidance can lead to additional problems to promote buy-in | • Revised quotations to be explicit about faculty realizing they could not avoid discussing problems and being glad they stopped avoiding<br>• Added text about what faculty would get out of participating in IMPACT (e.g., more productivity, lab reputation), with heavy emphasis in the initial/final modules |
| **Skills training comes too late:** Participants wanted to start learning techniques for talking about conflicts earlier in the intervention | • Consolidated modules and timing so that individuals are introduced to the skills training process starting at the beginning of Week 2<br>• Added more foreshadowing about later skills (e.g., edited quotations in Week 1 modules to include specific techniques that were discussed in later modules) |
| **Overly prescriptive advice:** Some modules had lists of steps, which seemed prescriptive rather than flexible and adaptable | • Avoided lists of numbered steps when possible<br>• Explicitly acknowledged that sets of steps could be adapted flexibly, giving examples of how that could look (e.g., in the “Discover” and “Decide” steps) |
| **Simplistic examples:** Participants wanted acknowledgement of the complexity of many real-life mentoring conflicts and student issues when examples seemed too “easy” | • Revised quotations and some examples so that issues were not always “solved” but instead improved somewhat over time as part of engaging in the outlined process<br>• Added multiple acknowledgements that real life scenarios may be more complex than what is depicted in the modules |
| **Wordiness:** Participants wanted interactive tasks, short tasks, bullet points, and engaging activities wherever possible | • Continually edited to make activities interactive, to shorten text where possible, and to add bullet points and quizzes/objects to click on instead of text blocks |

### Intervention Evaluation

After iteratively testing and revising the intervention materials, we conducted a field test of the IMPACT intervention. The design and methods for the field test are described below.

#### Participants

Enrollment in the field test took place in summer 2024. Individuals were eligible to participate if they were faculty members in a life science discipline (e.g., molecular biology, ecology, neuroscience) and had served as a research advisor to at least one PhD student for a minimum of one year. We direct emailed faculty at PhD-granting universities across the US about the study. We also contacted Directors of National Institutes of Health T32 Programs and asked them to share study information with faculty who met the eligibility criteria. We used a modified tailored panel management recruitment approach to encourage participation (Estrada et al., 2014), including providing compensation ($100 for the self-guided condition and $200 for the self-guided + peer discussion condition), emphasizing study and investigator credibility, and personalizing emails. The recruitment email included a link to an enrollment and screening survey, where faculty consented to participate and reported basic demographic information. Of the 86 faculty who started the enrollment survey, 8 did not consent and 7 did not complete the survey. A total of 71 eligible participants from 43 universities consented and constituted the sample for the field trial. Participant demographics are shown in Table 2, and a CONSORT diagram is provided in Figure 2.

**Figure 2.**
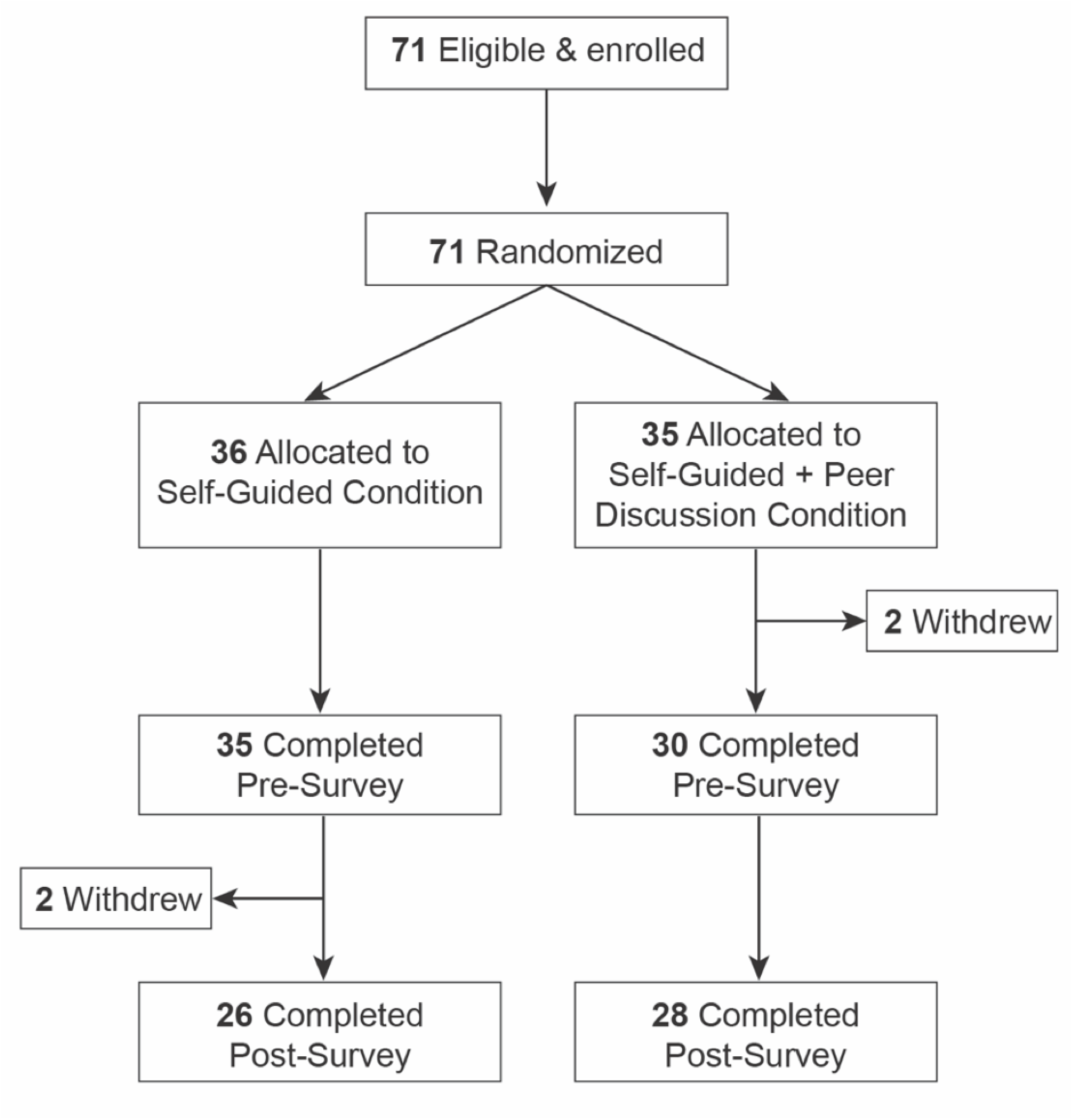
Participant flow diagram illustrating enrollment, randomization, and follow-up in accordance with CONSORT guidelines.

**Table 2.** Characteristics of faculty in field testing experiment (*n* = 71).

| Description | *n* (%) | Description | *n* (%) |
| --- | --- | --- | --- |
| **Gender** |  | **Years experience advising doctoral students** |  |
| Woman | 41 (58%) | 1 – 3 years | 17 (24%) |
| Man | 27 (38%) | 4 - 6 years | 15 (21%) |
| Non-binary | 1 (1%) | 7 - 10 years | 13 (18%) |
| Prefer not to respond | 2 (3%) | 11 - 15 years | 12 (17%) |
| **Race<sup>1</sup>** |  | 16 or more years | 14 (20%) |
| American Indian or Alaskan Native | 3 (4%) | **Current doctoral student advisees** |  |
| Asian | 6 (8%) | 1 | 16 (23%) |
| Black or African American | 2 (3%) | 2 | 21 (30%) |
| Hawaiian or Native Pacific Islander | 1 (1%) | 3 | 16 (23%) |
| Middle Eastern | 3 (4%) | 4 | 11 (15%) |
| White | 58 (82%) | 5 | 2 (3%) |
| Prefer to self-describe | 1 (1%) | 6 | 2 (3%) |
| Prefer not to respond | 3 (4%) | 7 or more | 2 (3%) |
| **Ethnicity** |  | **Prior participation in mentoring professional development** |  |
| Hispanic | 2 (3%) | Yes | 38 (54%) |
| Not Hispanic | 66 (93%) | No | 29 (41%) |
| Prefer not to respond | 3 (4%) | Unsure | 3 (4%) |
| **College generation status** |  | Prefer not to respond | 1 (1%) |
| First generation | 13 (18%) | **Total hours on mentor professional development<sup>2</sup>** |  |
| Continuing generation | 58 (82%) | 1 - 3 hours | 1 (3%) |
| **Rank** |  | 4 - 6 hours | 12 (41%) |
| Assistant Professor | 20 (28%) | 7 - 10 hours | 5 (13%) |
| Associate Professor | 27 (38%) | 11 - 15 hours | 8 (21%) |
| Full Professor | 24 (34%) | 16 or more hours | 14 (37%) |
| **Years as faculty** |  | Not sure | 1 (3%) |
| 1 - 3 years | 10 (14%) |  |  |
| 4 - 6 years | 9 (13%) |  |  |
| 7 - 10 years | 20 (28%) |  |  |
| 11 - 15 years | 12 (17%) |  |  |
| 16 or more years | 20 (28%) |  |  |

1 = Counts will not sum to 100% because participants could identify with multiple races.
2 = Percentages were calculated based on the subset of faculty participants who indicated they had engaged in some prior form of mentoring professional development.

#### Intervention Procedures

Participants were randomized to either the *Self-Guided Condition* or the *Self-Guided + Peer Discussion Condition.* To balance participants’ sociodemographic characteristics between the two conditions, we created four blocking groups based on: (1) gender (i.e., woman versus man and non-binary) and (2) career stage (i.e., pre-tenure/Assistant Professor versus post-tenure/Associate or Full Professor). Within each group, participants were randomly sorted and alternately assigned to the *Self-Guided Condition* (50%) or the *Self-Guided + Peer Discussion Condition* (50%). Participants were made aware of their condition upon assignment.

##### Implementation

As depicted in Figure 3, participants completed a pre-survey where they reported a) their attributions regarding the cause of conflicts with their graduate students, b) their self-efficacy for addressing conflict with graduate students, c) the quality of their mentoring relationship, d) their approaches for managing conflict, and e) the communication strategies and behaviors they engage in during conflicts, as well as their demographic information. Starting one week after the pre-survey, participants completed the 10 self-guided modules hosted on Qualtrics over a 6-week period, completing 1-2 modules per week depending on module length. The pacing was set by the research team (i.e., participants were only able to access modules during the designated week). Participants in the *Self-Guided + Peer Discussion Condition* also engaged in six 60-minute discussions on Zoom, which were facilitated by a subset of the research team and aligned with the weekly content of the modules. One week after completing the intervention, participants completed a post-survey, which assessed a) their attributions regarding the cause of conflicts with their graduate students, b) their self-efficacy for addressing conflict with graduate students, c) the quality of their mentoring relationship, d) their approaches for managing conflict, and e) their communication strategies and behaviors during conflicts.

**Figure 3.**
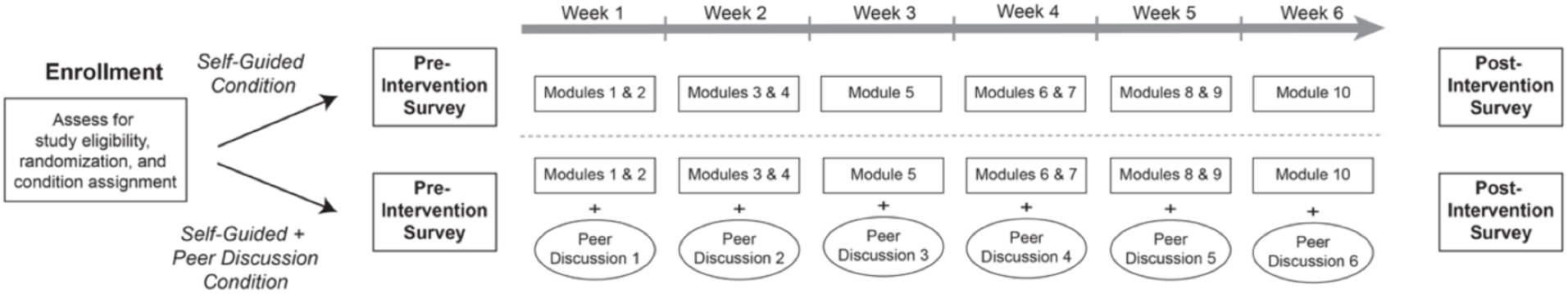
Study design for the IMPACT field test. Participants were randomized to the *Self-Guided Condition* or the *Self-Guided + Peer Discussion Condition.* Participations completed a pre-survey for baseline measures. One week later, all participants began the self-guided modules, which they completed over six weeks, with participants completing one or two modules per week. Participants in the *Self-Guided + Peer Discussion Condition* also participated in six weekly, 60-minute peer discussions on Zoom. One week after completing Week 6 activities, all participants completed a post-survey assessing the same variables as in the pre-survey.

#### Data Collection

Survey data were collected pre- and post-intervention via Qualtrics as depicted in Figure 3. Key outcomes of interest were motivational beliefs (i.e., self-efficacy to address conflicts and personal controllability attributions), engagement in conflict skills (i.e., collaborative versus avoidant conflict management, collaborative versus adversarial conflict communication, and engagement in specific practices from the IMPACT intervention), experiences with conflict (i.e., frequency and impacts of experienced conflict), and mentoring relationship satisfaction. A complete list of items is reported in the Supplemental Materials.

##### Self-efficacy

Self-efficacy to address conflicts with graduate students was assessed using a three-item scale adapted from Pintrich (1991) (*α*_pre_ = 0.80, *α*_post_ = 0.86). Responses were recorded on a five-point scale (i.e., 1 = “Not at all true”; 5 = “Completely true”). A sample item is *“I believe I can address conflicts that I experience with graduate students.”*

##### Personal controllability attributions

Personal control attributions regarding the causes of conflicts with graduate students were assessed using an attribution scale adapted from Coffee and Rees (2008). Participants were instructed to think about instances when they experienced conflicts with their graduate students and the *causes* of these conflicts. They then responded to three items measuring the extent to which they believed they had personal control over the causes of such conflicts (*α*_pre_ = 0.84, *α*_post_ = 0.85). Responses were recorded on a five-point scale (i.e., 1 = “Not at all”; 5 = “Completely”). A sample item is *“To what extent is the cause of conflicts with a graduate student something that you could influence in the future?”*

##### Experiences with conflict

Participants indicated the frequency and perceived impacts of the conflicts that occurred within their mentoring relationships with graduate students using five items developed for the present study. Responses were recorded on five-point frequency or extent scales (i.e., 1 = “Less than once a year/very little”; 5 = “More than once a week/an extensive amount”). Sample items included “*How often do you experience conflicts with your graduate students?”* and “*To what extent do conflicts with your graduate students impact your research productivity?”*

##### Collaborative conflict management

Behaviors related to finding mutually satisfactory solutions during conflict were assessed using a four-item scale adapted from Gelfand and colleagues (2012) (*α*_pre_ = 0.88, *α*_post_ = 0.71). Responses were recorded on a five-point scale (i.e., 1 = “Almost never”; 5 = “Almost every time”). A sample item is *“I examine issues with my graduate students until we find a solution that satisfies everyone.”*

##### Avoidant conflict management

Behaviors signaling a tendency to avoid open discussions with graduate students about conflicts were assessed using a four-item scale adapted from Gelfand and colleagues (2012) (*α*_pre_ = 0.75, *α*_post_ = 0.77). Responses were recorded on a five-point scale (i.e., 1 = “Almost never”; 5 = “Almost every time”). A sample item is *“I am very reluctant to openly talk about conflict.”*

##### Collaborative engagement in conflict communication

Behaviors reflective of constructive and prosocial communication during conflicts with graduate students (e.g., acknowledging the student’s perspective, communicating care or support, sharing thoughts in a constructive manner) were assessed using a seven-item scale adapted from Sanford (2010) (*α*_pre_ = 0.75, *α*_post_ = 0.70). Responses were recorded on a five-point scale (i.e., 1 = “Almost never”; 5 = “Almost every time”). A sample item is “*I make my students feel that their viewpoints are valuable.”*

##### Adversarial engagement in conflict communication

Behaviors reflecting critical, hostile, or defensive communication during conflicts with graduate students (e.g., voicing negative attributions, expressing contempt, criticism, or defensiveness) were assessed using an eight-item scale adapted from Sanford (2010) (*α*_pre_ = 0.70, *α*_post_ = 0.66). Responses were recorded on a five-point scale (i.e., 1 = “Almost never”; 5 = “Almost every time”). A sample item is “*I tell my students how they are doing something to cause the problem.”*

##### IMPACT-specific behaviors

Participants indicated how often they engaged in 15 specific behaviors that reflected the skills taught in IMPACT during their interactions with their graduate students. Responses were recorded on a five-point frequency scale (i.e., 1 = “Almost never”; 5 = “Almost every time”). Sample items included “*Asked open-ended questions to figure out your student’s perspective on the issue*” and “*Followed up to discuss whether the plan to address the issue was working*.”

##### Mentoring relationship satisfaction

Participants indicated their satisfaction with their mentoring relationship(s) with their graduate students(s) on a five-item scale from Allen and Eby (2003) (*α*_pre_ = 0.87, *α*_post_ = 0.88). Responses were recorded on a five-point scale (i.e., 1 = “Strongly disagree”; 5 = “Strongly agree”). A sample item is *“I am very satisfied with the mentoring relationship that my graduate students and I have developed.”*

#### Data Analysis

Our primary research question was how IMPACT participants’ motivational beliefs, self-reported experiences and behaviors related to conflict, and mentoring relationship satisfaction changed from pre- to post-intervention. Accordingly, we conducted paired-sample *t*-tests comparing pre-and post-scores for each measure, whether derived from multiple items or single-item scales, in *R* v4.1.0 in Rstudio (version 2022.12.353).

## RESULTS

We first present results regarding the feasibility of implementing both the self-guided version and the self-guided + peer discussion versions of IMPACT. Then we present results examining how mentors’ conflict-related motivational beliefs, self-reported behaviors and experiences with conflicts, and satisfaction with their graduate mentoring relationships changed pre- to post-IMPACT. We analyzed data for participants in both conditions together to maximize statistical power and address our primary study goals.

### Most participants completed the IMPACT intervention as intended

To evaluate the feasibility of implementing the IMPACT intervention, we tracked module completion and peer discussion attendance. Of the 71 faculty who initially enrolled in the study, 65 completed the pre-survey and 54 completed both the pre- and post-surveys (26 *Self-Guided* and *28 Self-Guided + Peer Discussion*) (Figure 2). Of these 54, participants in the *Self-Guided Condition* completed an average of 85% of the modules (range 73-96%). Participants in the *Self-Guided + Peer Discussion Condition* completed an average of 94% of the modules (range 89-96%) and attended 84% of the peer discussions (range 71-93%). These completion and attendance rates suggest good program adherence (i.e., faculty completed the intervention as intended).

### Peer discussions were implemented with fidelity

To ensure that the peer discussion sessions were delivered as intended, we assessed program fidelity following best practices for treatment integrity and fidelity monitoring (Bellg et al., 2004). Specifically, both facilitators independently completed a structured checklist for each peer discussion to indicate whether each required topic was covered. The facilitators also noted whether additional topics were discussed, whether any deviations occurred, which components were implemented particularly effectively, and whether there were any suggestions for future improvement. Results indicated that 100% of the intended checklist items were covered during the sessions, with 100% agreement between facilitators.

### Motivational beliefs and some conflict approaches improved post-IMPACT

Overall, participants showed statistically significant gains pre- to post-IMPACT in their motivational beliefs related to addressing conflicts with PhD students (Table 3 and Figure 4). Participants’ self-efficacy for addressing conflicts increased significantly (*t*(53) = −5.75, *p* < 0.0001, *d* = 0.80), as did their attributions that conflicts with PhD students can be managed through their personal actions (*t*(52) = −3.20, *p* = 0.002, *d* = 0.51), reflecting moderate to large gains. Participants also reported significantly increased collaborative engagement in conflict communication pre- to post-IMPACT (*t*(53) = −2.15, *p* = 0.036, *d* = 0.26). Participant reports of avoidant conflict management (*p =* 0.056), and adversarial engagement in conflict communication (*p =* 0.066) decreased pre- to post-IMPACT, but these changes were small and nonsignificant. Finally, participants reported similar levels of collaborative conflict management pre- to post-IMPACT (*p =* 0.343). Collectively, these results suggest that IMPACT enhanced participants’ motivational beliefs and some but not all of their approaches to managing conflicts with PhD students.

**Figure 4.**
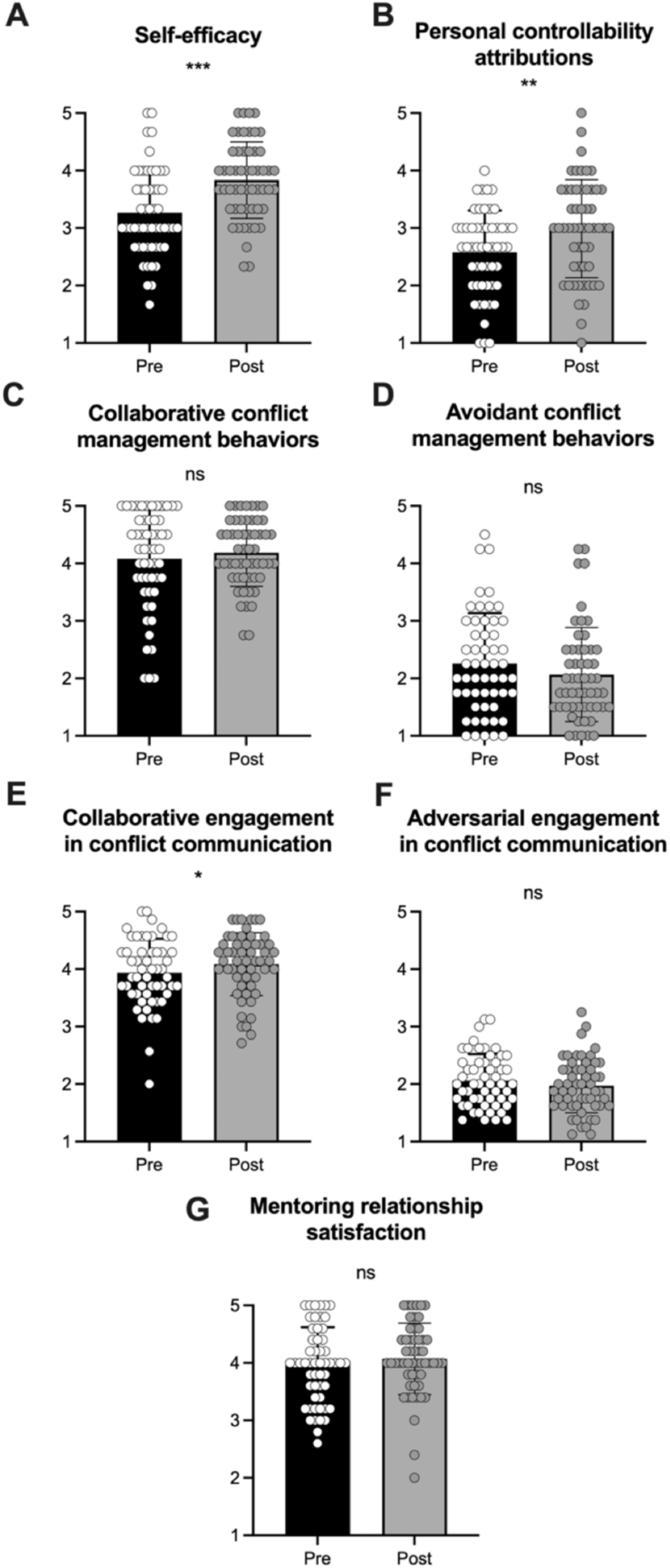
Results from the field test of IMPACT. *p-*values are indicated by asterisks: *\** < 0.05, ** < 0.01, *** < 0.001. ns = not significant.

**Table 3.**
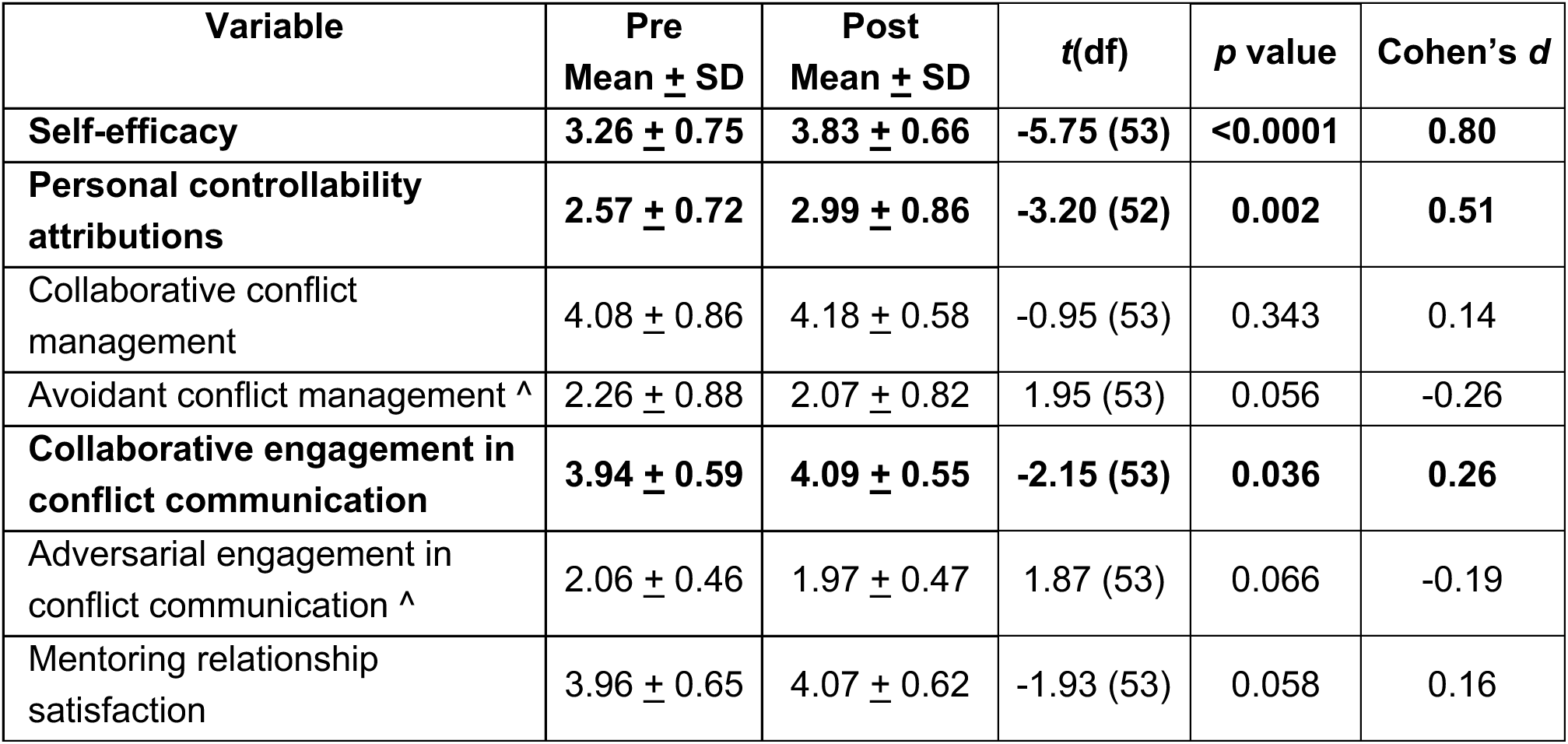
Pre- and post-IMPACT scores for all variables. Columns show variable values as means ± standard deviations. A caret (^) indicates variables for which the desired direction of the effect is negative (e.g., reduction in unproductive approach to addressing conflict). *t*(df) represents paired-samples t-tests comparing pre- and post-IMPACT scores. *p* < 0.05 are considered statistically significant and indicated with bold font. Cohen’s *d* indicates effect size; positive values reflect increases from pre- to post-intervention and negative values reflect decreases from pre- to post-IMPACT.

| Variable | Pre<br>Mean ± SD | Post<br>Mean ± SD | $t(df)$ | $p$ value | Cohen's $d$ |
| --- | --- | --- | --- | --- | --- |
| **Self-efficacy** | **3.26 ± 0.75** | **3.83 ± 0.66** | **-5.75 (53)** | **<0.0001** | **0.80** |
| **Personal controllability attributions** | **2.57 ± 0.72** | **2.99 ± 0.86** | **-3.20 (52)** | **0.002** | **0.51** |
| Collaborative conflict management | 4.08 ± 0.86 | 4.18 ± 0.58 | -0.95 (53) | 0.343 | 0.14 |
| Avoidant conflict management ^ | 2.26 ± 0.88 | 2.07 ± 0.82 | 1.95 (53) | 0.056 | -0.26 |
| **Collaborative engagement in conflict communication** | **3.94 ± 0.59** | **4.09 ± 0.55** | **-2.15 (53)** | **0.036** | **0.26** |
| Adversarial engagement in conflict communication ^ | 2.06 ± 0.46 | 1.97 ± 0.47 | 1.87 (53) | 0.066 | -0.19 |
| Mentoring relationship satisfaction | 3.96 ± 0.65 | 4.07 ± 0.62 | -1.93 (53) | 0.058 | 0.16 |

### Some but not all negative effects of conflict were reduced post-IMPACT

Participants reported experiencing fewer overall conflicts post-IMPACT (*t*(52) = 2.55, *p* = 0.013, *d* = −0.35), reflecting a significant and moderate reduction from pre-IMPACT levels (Table 4). Time and effort spent resolving conflicts with PhD students also decreased pre- to post-IMPACT (*t*(52) = 2.51, *p* = 0.015, *d* = −0.34), as did participants’ perceived impacts of conflicts on their research productivity (*t*(53) = 2.04, *p* = 0.046, *d* = −0.23). Emotional drain from conflicts decreased as well, though this change was not significant (*p* = 0.070) and the extent to which participants found conflicts upsetting also decreased but non-significantly (*p* = 0.137). Overall, these results indicate that, from pre- to post-IMPACT, participants experienced fewer conflicts with their PhD students, and these conflicts required less time and had a less negative impact on their research productivity. Although some effects did not reach statistical significance, the patterns were consistent with the IMPACT’s goal of reducing the frequency and negative consequences of conflicts.

**Table 4.** Pre- and post-intervention scores for mentors’ experiences with conflict. Columns show item means ± standard deviations. A caret (^) indicates items for which the desired direction of the effect is negative (e.g., reduction in conflict frequency). *t*(df) represents paired-samples t-tests comparing pre- and post-IMPACT scores. *p* < 0.05 are considered statistically significant and indicated with bold font. Cohen’s *d* indicates effect size; negative values reflect decreases from pre- to post-intervention.

| Variable | Pre<br>Mean ± SD | Post<br>Mean ± SD | $t(df)$ | $p$ value | Cohen's $d$ |
| --- | --- | --- | --- | --- | --- |
| How often do you experience conflicts with your graduate students? ^ | **2.17 ± 1.28** | **1.77 ± 0.85** | **2.55 (52)** | **0.013** | **-0.35** |
| How much time and effort do you typically spend resolving conflicts with your graduate students? ^ | **2.92 ± 1.27** | **2.53 ± 1.07** | **2.51 (52)** | **0.015** | **-0.34** |
| To what extent do conflicts with your graduate students leave you feeling emotionally drained? ^ | 3.35 ± 1.33 | 3.07 ± 1.26 | 1.84 (53) | 0.070 | -0.21 |
| To what extent do conflicts with your graduate students impact your research productivity? ^ | **2.93 ± 1.26** | **2.65 ± 1.17** | **2.04 (53)** | **0.046** | **-0.23** |
| To what extent do conflicts with your graduate students upset you? ^ | 3.37 ± 1.29 | 3.11 ± 1.16 | 1.51 (53) | 0.137 | -0.21 |

### Some but not all program-specific behaviors improved post-IMPACT

Participants reported significant improvements in many of the target behaviors emphasized in IMPACT from pre- to post-program (Table 5). Specifically, participants were less likely to bring up multiple issues they were concerned about during the same conversation (*t*(52) = 2.63, *p* = 0.011, *d* = −0.40); less likely to immediately suggest solutions to the issue (*t*(53) = 5.21, *p* < 0.001, d = −0.81); and less likely to tell students to calm down when their student got upset (*t*(53) = 2.41, *p* = 0.019, *d* = −0.37). Participants were also more likely to document a plan in writing about how they and their student would address the issue in the coming weeks (*t*(51) = - 2.46, *p* = 0.017, *d* = 0.44). These changes are consistent with the idea that IMPACT promoted many of the specific behaviors it was designed to address. Other target behaviors trended in the intended direction but did not reach statistical significance.

**Table 5.**
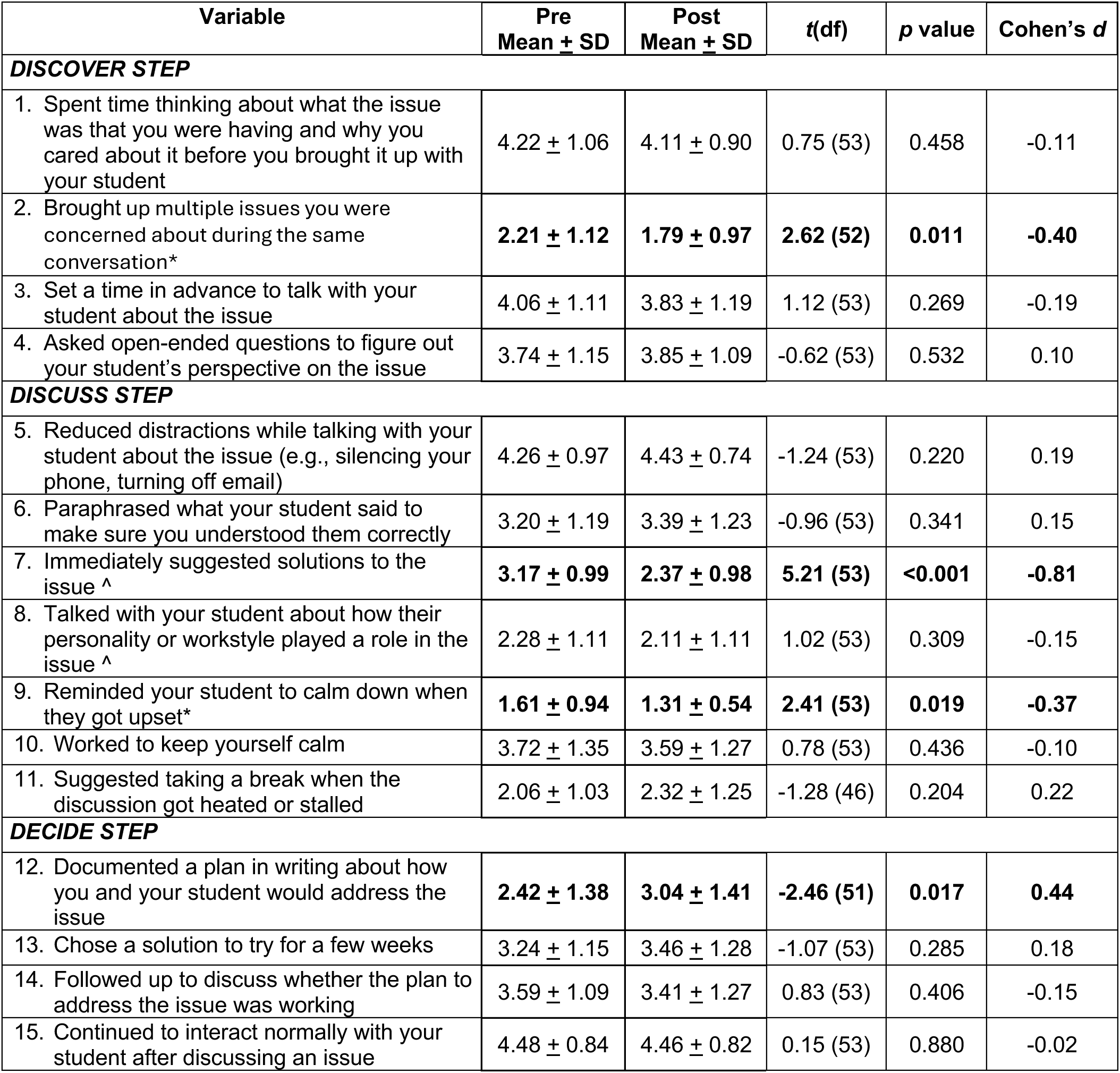
Pre- and post-IMPACT scores for IMPACT-specific behaviors. . Items represent behaviors related to each step of the three-step process for addressing conflicts with graduate students: Discover, Discuss, and Decide. A caret (^) indicates items for which the desired direction of the effect is negative (e.g., reduction in unproductive behavior). Columns show item means ± standard deviations. *t*(df) represents paired-samples t-tests comparing pre- and post-program scores. *p* < 0.05 is considered statistically significant and indicated with bold font. Cohen’s *d* indicates effect size.

| Variable | Pre Mean ± SD | Post Mean ± SD | $t(df)$ | $p$ value | Cohen's $d$ |
| --- | --- | --- | --- | --- | --- |
| **DISCOVER STEP** |  |  |  |  |  |
| 1. Spent time thinking about what the issue was that you were having and why you cared about it before you brought it up with your student | 4.22 ± 1.06 | 4.11 ± 0.90 | 0.75 (53) | 0.458 | -0.11 |
| 2. Brought up multiple issues you were concerned about during the same conversation* | **2.21 ± 1.12** | **1.79 ± 0.97** | **2.62 (52)** | **0.011** | **-0.40** |
| 3. Set a time in advance to talk with your student about the issue | 4.06 ± 1.11 | 3.83 ± 1.19 | 1.12 (53) | 0.269 | -0.19 |
| 4. Asked open-ended questions to figure out your student's perspective on the issue | 3.74 ± 1.15 | 3.85 ± 1.09 | -0.62 (53) | 0.532 | 0.10 |
| **DISCUSS STEP** |  |  |  |  |  |
| 5. Reduced distractions while talking with your student about the issue (e.g., silencing your phone, turning off email) | 4.26 ± 0.97 | 4.43 ± 0.74 | -1.24 (53) | 0.220 | 0.19 |
| 6. Paraphrased what your student said to make sure you understood them correctly | 3.20 ± 1.19 | 3.39 ± 1.23 | -0.96 (53) | 0.341 | 0.15 |
| 7. Immediately suggested solutions to the issue ^ | **3.17 ± 0.99** | **2.37 ± 0.98** | **5.21 (53)** | **<0.001** | **-0.81** |
| 8. Talked with your student about how their personality or workstyle played a role in the issue ^ | 2.28 ± 1.11 | 2.11 ± 1.11 | 1.02 (53) | 0.309 | -0.15 |
| 9. Reminded your student to calm down when they got upset* | **1.61 ± 0.94** | **1.31 ± 0.54** | **2.41 (53)** | **0.019** | **-0.37** |
| 10. Worked to keep yourself calm | 3.72 ± 1.35 | 3.59 ± 1.27 | 0.78 (53) | 0.436 | -0.10 |
| 11. Suggested taking a break when the discussion got heated or stalled | 2.06 ± 1.03 | 2.32 ± 1.25 | -1.28 (46) | 0.204 | 0.22 |
| **DECIDE STEP** |  |  |  |  |  |
| 12. Documented a plan in writing about how you and your student would address the issue | **2.42 ± 1.38** | **3.04 ± 1.41** | **-2.46 (51)** | **0.017** | **0.44** |
| 13. Chose a solution to try for a few weeks | 3.24 ± 1.15 | 3.46 ± 1.28 | -1.07 (53) | 0.285 | 0.18 |
| 14. Followed up to discuss whether the plan to address the issue was working | 3.59 ± 1.09 | 3.41 ± 1.27 | 0.83 (53) | 0.406 | -0.15 |
| 15. Continued to interact normally with your student after discussing an issue | 4.48 ± 0.84 | 4.46 ± 0.82 | 0.15 (53) | 0.880 | -0.02 |

### Mentoring relationship satisfaction was unchanged post-IMPACT

Although participants reported greater satisfaction with their mentoring relationships with graduate students from pre- to post-IMPACT, this effect was small and did not reach statistical significance (*p =* 0.058) (Table 3).

#### Exploratory supplemental analyses

We combined data from participants in both conditions for the analyses described above. As a final step, we conducted exploratory supplemental analyses to assess potential differences between the *Self-Guided* Condition and *Self-Guided + Peer Discussion* Condition. Overall, the independent-samples *t*-tests indicated that participants in both conditions were comparable at the pre-intervention measure, with all but one construct showing no significant between-group differences (all *p* values > 0.05) and only a few items showing between-group differences. The results of the analyses are reported in the Supplemental Materials.

Although the study was not powered to detect between-condition effects, we conducted two complementary sets of analyses to examine the extent to which the two versions of the intervention yielded distinct patterns of change at the post-intervention measure. For the items and outcomes that showed no significant baseline differences (α ≥ 0.05) between the *Self-Guided* Condition and *Self-Guided + Peer Discussion* Condition at the pre-intervention measure, we conducted independent-samples t-tests to compare post-intervention scores. For items and outcomes that did show significant differences between groups at the pre-intervention measure, we conducted analyses of covariance (ANCOVAs) with post-intervention scores as the dependent variable, group as the between-subjects factor, and baseline scores as a covariate. This approach allowed us to control for pre-existing differences while assessing the effect of program format on post-intervention outcomes. Our results suggest that the two conditions yielded similar post-intervention outcomes; however, these findings should be interpreted with caution due to the small sample sizes.

## DISCUSSION

This paper describes the development, refinement, and initial implementation of the IMPACT intervention, a mentoring professional development program for life science faculty that aims to foster their self-efficacy beliefs and build their skills so that they can navigate conflicts with their graduate students more effectively. IMPACT meets a critical need given that conflicts arise in graduate mentoring relationships and graduate students describe their mentors’ responses to these conflicts as influencing their graduate experiences and outcomes (Tuma et al., 2021). Furthermore, existing professional development efforts are not designed to help mentors improve their abilities to navigate conflicts (Gangrade et al., 2024). We described the process of designing IMPACT to illustrate how a theory- and principle-based approach can be applied to developing and refining an intervention. This work drew on interdisciplinary research on motivation (Graham & Taylor, 2022), wise interventions (Walton & Wilson, 2018; Yeager & Walton, 2011), interpersonal communication (Epstein & Baucom, 2002), and learning (Bandura & Walters, 1977; Fiorella & Mayer, 2016; Lave & Wenger, 1991), while aligning with the norms and culture of academic life science research. By detailing our methodological choices and how feedback was incorporated to iteratively refine the intervention materials, we provide a transparent example for others designing theory-informed interventions.

For an intervention to be practically effective, it must be feasible to implement and useful to the target audience. Through our field test, we found that IMPACT was feasible to implement with fidelity, and mentors were willing to complete IMPACT at a high rate, despite the time demands. However, this high engagement may reflect selection bias, as faculty volunteered to participate and may be more committed to completing the program than a broader population of life science faculty. The delivery modes (online, asynchronous) and platforms (Qualtrics, Zoom) may have contributed to feasibility and acceptability because mentors could complete the modules at a time that worked for them and could participate in peer discussions from any location. In addition, delivery of the peer discussions over Zoom supported fidelity of implementation because facilitators could consult structured guidance to ensure consistent delivery and emphasis of key messages. These features of IMPACT increase its potential to be transferable and scalable.

Field test participants reported improved beliefs about their ability to address conflicts with their graduate students and improved skills related to addressing conflicts productively, including the use of IMPACT-specific behaviors, following the intervention. They also reported fewer conflicts with their PhD students and that these conflicts required less of their time and had a less negative impact on their research productivity. Collectively, these results suggest that IMPACT is successfully changing both intended mechanisms (i.e., motivation, skill-building), and that addressing *both* beliefs *and* skills likely contributes to IMPACT’s effectiveness. These results are exciting because they speak to the power of a theory-informed intervention aimed at addressing conflicts in mentoring by targeting both the beliefs that underlie skill-building and key skills themselves. By targeting both beliefs and skills, using effective techniques and strategies from prior work, the IMPACT intervention produced robust benefits while maintaining feasibility. Another benefit of IMPACT’s design is that it is likely to help mentors apply conflict management skills beyond the intervention period, thereby potentially reducing the frequency and impacts of conflicts in the future. Because the intervention tapped into recursive psychological processes that underlie behavior change, while building skills to actually change behavior (i.e., beliefs about engaging in conflict management, development of conflict management skills, and conflict outcomes affect one another cyclically over time), mentors may continue benefitting from these belief-behavior patterns into the future. Longer-term and longitudinal research is necessary to yield insight into how mentors’ conflict-related beliefs, skills, and behaviors evolve over time and to determine the long-term impact of these changes on mentors and graduate students. However, the potential for this intervention to build skills longer-term is high, given that research on wise interventions, in which IMPACT was grounded, indicates that positive effects can continue to accrue for months and even years (Walton & Wilson, 2018; Yeager & Walton, 2011).

Collectively, these results are promising and support further research on IMPACT, including evaluation in a randomized controlled trial (RCT) with a no-treatment control condition. Such an experimental design would allow for causal conclusions about IMPACT’s effects. If future research shows that mentors who participate in IMPACT are more motivated and skilled to address conflicts with their graduate students, this could favorably influence graduate student mentoring and improve the overall quality of the graduate education experience. In addition, this type of design would allow us to compare both types of treatment conditions separately to a no-treatment control condition. If the *Self-Guided* version is sufficient to improve mentors’ motivational beliefs and conflict management skills compared with a control condition, such evidence could justify its wider dissemination. Importantly, *Self-Guided* IMPACT requires ∼6 hours of faculty time, whereas ∼12 hours are needed for the *Self-Guided + Peer Discussion* version. In addition, implementing the *Self-Guided + Peer Discussion* version requires skilled facilitation and greater coordination (i.e., ensuring participants complete self-guided materials prior to the peer discussion each week). Testing both versions in a sufficiently powered study would yield insight into whether participation in the facilitated peer discussion is needed to maximize the benefits of IMPACT.

Future research on IMPACT effectiveness should also include graduate students’ perspectives to complement mentor reports. We opted to focus on collecting data only from mentors for this initial field test given the time and effort required to collect dyadic data. Yet self-report is a limited measure of skills. Mentors may be rating their skills more highly after IMPACT even though their actual skills may be unchanged, either because they want to appear knowledgeable of best practices or because they are not accurately perceiving their own behavior (Kruger & Dunning, 1999). Collecting reports from both faculty and their graduate students would allow for more valid inferences about IMPACT’s effectiveness and contribute to the very limited body of knowledge about dyadic perspectives on mentoring (Allen et al., 2008; Hernandez, 2018). Examining the experiences of graduate students whose mentors complete IMPACT (or not) would also allow for more complete understanding of the program’s benefits.

In sum, the IMPACT intervention combined attribution retraining and conflict management skill building to help life science faculty members better manage conflict with their graduate students. Promising field test results of this theory-based, recursively designed intervention support further testing to better understand potential impacts on life science faculty and their graduate students.

## ACKNOWLEDGMENTS

This work was supported in part by funding from the Georgia Athletic Association Professorship for Innovative Science Education, National Institute of General Medical Sciences of the National Institutes of Health Award Number R01GM147061, and National Science Foundation Division of Graduate Education Grant Number 2328692. The content is solely the responsibility of the authors and does not necessarily represent the official views of the National Institutes of Health or the National Science Foundation.

## Footnotes

1 We will use the term “mentor” when referring to the principal investigator or faculty member who serves as primary advisor for a graduate student’s dissertation research from here forward.

