## Supplemental materials for "Development and field test of an intervention to reduce conflict in faculty-doctoral student mentoring relationships"

**Table of Contents**

| Contents | Pages |
|---|---|
| Measures | 2-5 |
| Exploratory Supplemental Analyses | 6-12 |

### MEASURES

#### Personal controllability attributions

Coffee, P., & Rees, T. (2008). The CSGU: A measure of controllability, stability, globality, and universality attributions. *Journal of Sport and Exercise Psychology,* 30(5), 611–641.

**Instructions:** Think about times when you experience conflicts with your graduate students and the **causes** of these conflicts. Please indicate the extent to which you think the causes of these conflicts are something that...
**Response Scale:** 1 = Not at all, 2 = A little, 3 = Somewhat, 4 = A lot, 5 = Completely

| |
|---|
| 1. You could influence in the future. |
| 2. You could alter in the future. |
| 3. You could control in the future. |

#### Self-efficacy

Pintrich, P. R. (1991). *A manual for the use of the Motivated Strategies for Learning Questionnaire (MSLQ).* https://eric.ed.gov/?id=ED338122

**Instructions:** The next set of statements asks about how you view conflicts with your graduate students and your ability to navigate them. Please rate how true the following statements are.
**Response Scale:** 1 = Not at all true, 2 = A little true, 3 = Somewhat true, 4 = A lot true, 5 = Completely true

| |
|---|
| 1. I believe I can address conflicts that I experience with my graduate students. |
| 2. I'm confident I can address basic conflicts that I experience with my graduate students. |
| 3. I'm certain I can address the most challenging conflicts that I experience with my graduate students. |

#### Mentoring relationship satisfaction

Allen, T. D., & Eby, L. T. (2003). Relationship effectiveness for mentors: Factors associated with learning and quality. *Journal of Management*, *29*(4), 469–486. https://doi.org/10.1016/S0149-2063_03_00021-7

**Instructions:** Please indicate the extent to which you agree with the following statements.
**Response Scale:** 1 = Strongly disagree, 2 = Disagree, 3 = Undecided, 4 = Agree, 5 = Strongly agree

| |
|---|
| 1. The relationship I have with my graduate students is very effective. |
| 2. I am very satisfied with the mentoring relationship that my graduate students and I have developed. |
| 3. I am effectively utilized as a mentor by my graduate students. |
| 4. My graduate students and I enjoy a high-quality relationship. |
| 5. Both my graduate students and I benefit from our mentoring relationship. |

**Collaborative conflict management**
Gelfand, M. J., Leslie, L. M., Keller, K., & de Dreu, C. (2012). Conflict cultures in organizations: How leaders shape conflict cultures and their organizational-level consequences. *Journal of Applied Psychology*, *97*(6), 1131-1147.

**Instructions:** Please indicate the extent to which you do the following.
**Response Scale:** 1 = Almost never, 2 = Sometimes, 3 = About half the time, 4 = Often, 5 = Almost every time

| |
|---|
| 1. I examine issues with my graduate students until we find a solution that satisfies everyone. |
| 2. I examine ideas from all sides to find a mutually optimal solution. |
| 3. I try my best to work out a solution with my graduate students that serves everyone's interests. |
| 4. I try to come up with creative solutions that incorporate multiple perspectives. |

**Avoidant conflict management**
Gelfand, M. J., Leslie, L. M., Keller, K., & de Dreu, C. (2012). Conflict cultures in organizations: How leaders shape conflict cultures and their organizational-level consequences. *Journal of Applied Psychology*, *97*(6), 1131-1147.

**Instructions:** Please indicate the extent to which you do the following.
**Response Scale:** 1 = Almost never, 2 = Sometimes, 3 = About half the time, 4 = Often, 5 = Almost every time

| |
|---|
| 1. I discuss conflict with my graduate students openly. |
| 2. I avoid openly discussing conflicts with my graduate students. |
| 3. I am very reluctant to openly talk about conflict. |
| 4. Conflict is dealt with openly in my lab. |

**Collaborative engagement in conflict communication**
Sanford, K. (2010). Assessing conflict communication in couples: Comparing the validity of self-report, partner-report, and observer ratings. *Journal of Family Psychology*, *24*(2), 165-174. https://doi.org/10.1037/a0017953

**Instructions:** Please indicate the extent to which you do the following during conflicts with your graduate students.
**Response Scale:** 1 = Almost never, 2 = Sometimes, 3 = About half the time, 4 = Often, 5 = Almost every time

| |
|---|
| 1. I make my students feel that their viewpoints are valuable. |
| 2. I say something kind. |
| 3. I am considerate toward my students. |
| 4. I agree with my students. |
| 5. I politely talk about my feelings. |
| 6. I carefully listen so I can understand my students. |
| 7. I discuss the issue calmly. |

**Adversarial engagement in conflict communication**

Sanford, K. (2010). Assessing conflict communication in couples: Comparing the validity of self-report, partner-report, and observer ratings. *Journal of Family Psychology*, *24*(2), 165-174. https://doi.org/10.1037/a0017953.

**Instructions:** Please indicate the extent to which you do the following during conflicts with your graduate students.
**Response Scale:** 1 = Almost never, 2 = Sometimes, 3 = About half the time, 4 = Often, 5 = Almost every time

| |
|---|
| 1. I raise my voice. |
| 2. I tell my students how they are doing something to cause the problem. |
| 3. I say something mean. |
| 4. I argue. |
| 5. I defend my position. |
| 6. I correct my students' statements that are not true. |
| 7. I criticize my students. |
| 8. I tell my students to calm down. |

**Experiences with conflict**

Investigator-generated.

**Instructions:** Conflicts are a part of any long-term relationship. Since we have long-term relationships with our graduate students, it is likely we will experience conflicts at some point.
**Response Scale for item 1:** 1 = Less than once a year, 2 = Once a year, 3 = Once every few months, 4 = Once a month, 5 = Once a week, 6 = More than once a week
**Response Scale for items 2-5:** 1 = Very little, 2 = A little, 3 = A moderate amount, 4 = A good amount, 5 = An extensive amount

| |
|---|
| 1. How often do you experience conflicts with your graduate students? |
| 2. How much time and effort do you typically spend resolving conflicts with your graduate students? |
| 3. To what extent do conflicts with your graduate students leave you feeling emotionally drained? |
| 4. To what extent do conflicts with your graduate students impact your research productivity? |
| 5. To what extent do conflicts with your graduate students upset you? |

**IMPACT-specific behaviors**
Investigator-generated.

**Instructions:** Please rate how often you have done the following when you have experienced a conflict or disagreement regarding an issue with a graduate student.
**Response Scale:** 1 = Almost never, 2 = Sometimes, 3 = About half the time, 4 = Often, 5 = Almost every time

| |
|---|
| ***DISCOVER STEP*** |
| 1. Spent time thinking about what the issue was that you were having and why you cared about it before you brought it up with your student. |
| 2. Brought up multiple issues you were concerned about during the same conversation. |
| 3. Set a time in advance to talk with your student about the issue. |
| 4. Asked open-ended questions to figure out your student's perspective on the issue. |
| ***DISCUSS STEP*** |
| 5. Reduced distractions while talking with your student about the issue (e.g., silencing your phone, turning off email). |
| 6. Paraphrased what your student said to make sure you understood them correctly. |
| 7. Immediately suggested solutions to the issue. |
| 8. Talked with your student about how their personality or workstyle played a role in the issue. |
| 9. Reminded your student to calm down when they got upset. |
| 10. Worked to keep yourself calm. |
| 11. Suggested taking a break when the discussion got heated or stalled. |
| ***DECIDE STEP*** |
| 12. Documented a plan in writing about how you and your student would address the issue. |
| 13. Chose a solution to try for a few weeks. |
| 14. Followed up to discuss whether the plan to address the issue was working. |
| 15. Continued to interact normally with your student after discussing an issue. |

### EXPLORATORY SUPPLEMENTAL ANALYSIS

#### Pre-intervention differences between conditions

We conducted independent-samples t-tests to examine potential differences between the *Self-Guided* Condition and the *Self-Guided* + *Peer Discussion* Condition pre-intervention. Overall, our results indicate that the conditions were largely comparable at baseline, with most items and constructs showing no significant between-group differences (all *ps* > .05). Complete results are reported below.

**Cross-condition comparison of key variables pre-intervention.** $p < 0.05$ is considered statistically significant and indicated in bold.

| Variable | Self-Guided Condition Mean ± SD | Self-Guided + Peer Discussion Condition Mean ± SD | *t*(df) | *p* value |
|---|---|---|---|---|
| Self-efficacy | 3.29 ± 0.79 | 3.24 ± 0.73 | 0.28 (50.78) | 0.784 |
| Personal controllability attributions | 2.54 ± 0.74 | 2.62 ± 0.73 | -0.39 (50.85) | 0.696 |
| Collaborative conflict management | 4.23 ± 0.83 | 3.94 ± 0.87 | 1.26 (51.95) | 0.212 |
| Avoidant conflict management | 2.07 ± 0.80 | 2.44 ± 0.92 | -1.58 (51.76) | 0.120 |
| **Collaborative engagement in conflict communication** | **4.12 ± 0.50** | **3.77 ± 0.62** | **2.31 (51.02)** | **0.025** |
| Adversarial engagement in conflict communication | 2.00 ± 0.48 | 2.12 ± 0.44 | -0.92 (50.53) | 0.361 |
| Mentoring relationship satisfaction | 3.97 ± 0.68 | 3.96 ± 0.65 | 0.66 (51.32) | 0.947 |

**Cross-condition comparison of mentor experiences with conflict pre-intervention.** $p < 0.05$ is considered statistically significant and indicated in bold.

| **Variable** | **Self-Guided Condition Mean ± SD** | **Self-Guided + Peer Discussion Condition Mean ± SD** | ***t*(df)** | ***p* value** |
|---|---|---|---|---|
| How often do you experience conflicts with your graduate students? | 2.23 ± 1.37 | 2.11 ± 1.22 | 0.33 (49.86) | 0.738 |
| **How much time and effort do you typically spend resolving conflicts with your graduate students?** | **3.35 ± 1.26** | **2.52 ± 1.16** | **2.48 (50.19)** | **0.016** |
| To what extent do conflicts with your graduate students leave you feeling emotionally drained? | 3.54 ± 1.17 | 3.18 ± 1.47 | 0.99 (50.92) | 0.322 |
| To what extent do conflicts with your graduate students impact your research productivity? | 2.96 ± 1.34 | 2.89 ± 1.20 | 0.20 (50.22) | 0.843 |
| To what extent do conflicts with your graduate students upset you? | 3.42 ± 1.17 | 3.32 ± 1.42 | 0.29 (51.35) | 0.774 |

**Cross-condition comparison of IMPACT-specific behaviors pre-intervention.** *p* < 0.05 is considered statistically significant and indicated in bold.

| Variable | Self-Guided Condition Mean ± SD | Self-Guided + Peer Discussion Condition Mean ± SD | *t*(df) | *p* value |
|---|---|---|---|---|
| ***DISCOVER STEP*** | | | | |
| 1. Spent time thinking about what the issue was that you were having and why you cared about it before you brought it up with your student | 4.35 ± 1.13 | 4.11 ± 0.99 | 0.82 (49.97) | 0.414 |
| 2. Brought up multiple issues you were concerned about during the same conversation | 1.96 ± 1.00 | 2.44 ± 1.19 | -1.60 (50.11) | 0.115 |
| **3. Set a time in advance to talk with your student about the issue** | **4.42 ± 0.86** | **3.71 ± 1.21** | **2.49 (48.64)** | **0.016** |
| 4. Asked open-ended questions to figure out your student's perspective on the issue | 4.00 ± 1.02 | 3.50 ± 1.23 | 1.62 (51.34) | 0.109 |
| 5. Reduced distractions while talking with your student about the issue (e.g., silencing your phone, turning off email) | 4.35 ± 0.75 | 4.18 ± 1.16 | 0.637 (46.51) | 0.527 |
| 6. Paraphrased what your student said to make sure you understood them correctly | 3.38 ± 1.13 | 3.04 ± 1.23 | 1.08 (51.99) | 0.283 |
| 7. Immediately suggested solutions to the issue | 3.19 ± 0.98 | 3.14 ± 1.01 | 0.18 (51.88) | 0.855 |
| 8. Talked with your student about how their personality or workstyle played a role in the issue | 2.12 ± 1.11 | 2.43 ± 1.10 | -1.04 (51.67) | 0.303 |
| 9. Reminded your student to calm down when they got upset | 1.46 ± 0.76 | 1.75 ± 1.08 | -1.14 (48.66) | 0.258 |
| 10. Worked to keep yourself calm | 3.54 ± 1.50 | 3.89 ± 1.20 | -0.95 (47.77) | 0.344 |
| 11. Suggested taking a break when the discussion got heated or stalled | 2.00 ± 1.08 | 2.16 ± 1.14 | -0.51 (47.85) | 0.613 |
| ***DECIDE STEP*** | | | | |
| 12. Documented a plan in writing about how you and your student would address the issue | 2.46 ± 1.48 | 2.44 ± 1.31 | 0.04 (49.78) | 0.964 |
| 13. Chose a solution to try for a few weeks | 3.19 ± 1.17 | 3.29 ± 1.15 | -0.29 (51.58) | 0.769 |
| 14. Followed up to discuss whether the plan to address the issue was working | 3.73 ± 1.04 | 3.46 ± 1.14 | -0.89 (51.99) | 0.373 |
| 15. Continued to interact normally with your student after discussing an issue | 4.62 ± 0.70 | 4.36 ± 0.95 | 1.14 (49.42) | 0.258 |

**Post-intervention differences between conditions**

To examine potential post-intervention differences between the *Self-Guided* and *Self-Guided + Peer Discussion* Conditions, we conducted two complementary sets of analyses. We found no statistically significant post-intervention differences between the two conditions for any of the analyses.

*Items and variables without baseline differences*
For the items and variables that did not differ between the two conditions pre-intervention, we conducted independent-samples t-tests to compare post-intervention differences. The results are reported below.

**Cross-condition comparison of key variables post-intervention.**

| Variable | Self-Guided Only Mean $\pm$ SD | Self-Guided + Peer Discussion Mean $\pm$ SD | $t$(df) | $p$ value | Cohen's $d$ |
|---|---|---|---|---|---|
| Self-efficacy | 3.71 $\pm$ 0.66 | 3.95 $\pm$ 0.67 | -1.38 (51.80) | 0.175 | 0.37 |
| Personal controllability attributions | 2.86 $\pm$ 0.77 | 3.11 $\pm$ 0.92 | -1.07 (51.49) | 0.287 | 0.29 |
| Collaborative conflict management | 4.26 $\pm$ 0.49 | 4.12 $\pm$ 0.66 | 0.91 (49.52) | 0.366 | -0.25 |
| Avoidant conflict management | 2.04 $\pm$ 0.92 | 2.09 $\pm$ 0.73 | -0.23 (47.55) | 0.814 | 0.07 |
| Adversarial engagement in conflict communication | 1.86 $\pm$ 0.47 | 2.08 $\pm$ 0.46 | -1.71 (51.57) | 0.093 | 0.46 |
| Mentoring relationship satisfaction | 4.15 $\pm$ 0.54 | 3.99 $\pm$ 0.69 | 0.96 (50.80) | 0.342 | -0.26 |

**Cross-condition comparison of mentor experiences with conflict post-intervention.**

| Variable | Self-Guided Only Mean $\pm$ SD | Self-Guided + Peer Discussion Mean $\pm$ SD | $t$(df) | $p$ value | Cohen's $d$ |
|---|---|---|---|---|---|
| How often do you experience conflicts with your graduate students? | 1.58 $\pm$ 0.70 | 1.93 $\pm$ 0.94 | -1.15 (49.80) | 0.124 | -0.42 |
| To what extent do conflicts with your graduate students leave you feeling emotionally drained? | 3.08 $\pm$ 1.32 | 3.07 $\pm$ 1.21 | 0.02 (50.68) | 0.987 | 0.00 |
| To what extent do conflicts with your graduate students impact your research productivity? | 2.54 $\pm$ 1.24 | 2.75 $\pm$ 1.11 | -0.66 (50.26) | 0.513 | -0.18 |
| To what extent do conflicts with your graduate students upset you? | 3.19 $\pm$ 1.13 | 3.04 $\pm$ 1.20 | 0.49 (51.99) | 0.624 | 0.13 |

**Cross-condition comparison of IMPACT-specific behaviors post-intervention.**

| Variable | Self-Guided Only Mean ± SD | Self-Guided + Peer Discussion Mean ± SD | *t*(df) | *p* value | Cohen's *d* |
|---|---|---|---|---|---|
| ***DISCOVER STEP*** | | | | | |
| 1. Spent time thinking about what the issue was that you were having and why you cared about it before you brought it up with your student | 4.12 ± 0.86 | 4.11 ± 0.964 | 0.03 (51.97) | 0.974 | 0.01 |
| 2. Brought up multiple issues you were concerned about during the same conversation | 1.65 ± 0.98 | 1.89 ± 0.96 | -0.90 (51.51) | 0.369 | -0.24 |
| 4. Asked open-ended questions to figure out your student's perspective on the issue | 3.88 ± 0.99 | 3.82 ± 1.19 | 0.21 (51.45) | 0.832 | 0.06 |
| ***DISCUSS STEP*** | | | | | |
| 5. Reduced distractions while talking with your student about the issue (e.g., silencing your phone, turning off email) | 4.35 ± 0.74 | 4.50 ± 0.74 | -0.78 (51.71) | 0.452 | -0.21 |
| 6. Paraphrased what your student said to make sure you understood them correctly | 3.54 ± 1.14 | 3.25 ± 1.32 | 0.86 (51.72) | 0.393 | 0.23 |
| 7. Immediately suggested solutions to the issue | 2.23 ± 0.86 | 2.50 ± 1.07 | -1.02 (51.01) | 0.313 | -0.27 |
| 8. Talked with your student about how their personality or workstyle played a role in the issue | 2.23 ± 1.31 | 2.00 ± 0.90 | 0.75 (44.07) | 0.457 | 0.21 |
| 9. Reminded your student to calm down when they got upset | 1.27 ± 0.53 | 1.36 ± 0.56 | -0.59 (51.96) | 0.557 | -0.16 |
| 10. Worked to keep yourself calm | 3.77 ± 1.18 | 3.43 ± 1.35 | 0.99 (51.82) | 0.325 | 0.27 |
| 11. Suggested taking a break when the discussion got heated or stalled | 2.46 ± 1.22 | 2.31 ± 1.41 | 0.405 (47.80) | 0.686 | 0.11 |
| ***DECIDE STEP*** | | | | | |
| 12. Documented a plan in writing about how you and your student would address the issue | 3.16 ± 1.37 | 2.96 ± 1.45 | 0.503 (50.81) | 0.617 | 0.14 |
| 13. Chose a solution to try for a few weeks | 3.65 ± 1.02 | 3.29 ± 1.49 | 1.06 (47.93) | 0.291 | 0.29 |
| 14. Followed up to discuss whether the plan to address the issue was working | 3.54 ± 1.14 | 3.29 ± 1.38 | 0.734 (51.29) | 0.466 | 0.20 |
| 15. Continued to interact normally with your student after discussing an issue | 4.54 ± 0.58 | 4.39 ± 0.99 | 0.662 (44.11) | 0.511 | 0.18 |

*Items and variables with baseline differences*
For the one variable (i.e., collaborative engagement in conflict communication) and two items that differed between the two conditions pre-intervention, we conducted analyses of covariances (ANCOVAs) to examine potential differences post-intervention. Post-intervention scores were included as the dependent variable, condition as the between-subjects factor, and pre-intervention scores as a covariate. We calculated partial $\eta^2$ values to estimate effect sizes for each ANCOVA and estimated marginal means with 95% confidence intervals to facilitate the interpretation of adjusted post-intervention differences by condition.

*1. Collaborative engagement in conflict communication*
After controlling for pre-intervention levels, there was no significant effect of condition, $F(1,51) = 0.0004$, $p = 0.984$, partial $\eta^2 = 0.05$, on post-intervention levels.

**ANCOVA results**

| | **df** | **F** | ***p*** | **Partial $\eta^2$** |
|---|---|---|---|---|
| Condition (*Self-Guided* vs. *Self-Guided + Peer Discussion*) | 1 | 0.0004 | 0.984 | 0.05 |
| Pre-intervention collaborative engagement in conflict communication | 1 | 26.75 | <0.001 | 0.34 |
| Residual | 51 | | | |

**Estimated marginal means**

| **Condition** | **Adjusted Mean** | **SE** | **95% CI** |
|---|---|---|---|
| *Self-Guided* | 4.08 | 0.09 | [3.90, 4.26] |
| *Self-Guided + Peer Discussion* | 4.09 | 0.09 | [3.91, 4.26] |

*2. How much time and effort do you typically spend resolving conflicts with your graduate students?*
After controlling for pre-intervention levels, there was no significant effect of condition, $F(1,50) = 1.31$, $p = 0.257$, partial $\eta^2 = 0.003$, on post-intervention levels.

**ANCOVA results**

| | **df** | **F** | ***p*** | **Partial $\eta^2$** |
|---|---|---|---|---|
| Condition (*Self-Guided* vs. *Self-Guided + Peer Discussion* ) | 1 | 1.31 | 0.257 | 0.003 |
| Pre-intervention time and effort | 1 | 20.92 | <0.001 | 0.30 |
| Residual | 50 | | | |

**Estimated marginal means**

| **Condition** | **Adjusted Mean** | **SE** | **95% CI** |
|---|---|---|---|
| *Self-Guided* | 2.37 | 0.18 | [2.00, 2.74] |
| *Self-Guided + Peer Discussion* | 2.68 | 0.18 | [2.31, 3.04] |

3. *Set a time in advance to talk with your student about the issue*
After controlling for pre-intervention levels, there was no significant effect of condition, $F(1,51) = 2.45$, $p = 0.123$, partial $\eta^2 = 0.07$, on post-intervention levels.

**ANCOVA results**

| | df | F | *p* | Partial $\eta^2$ |
|---|---|---|---|---|
| Condition (*Self-Guided* vs. *Self-Guided + Peer Discussion*) | 1 | 2.45 | 0.123 | 0.07 |
| Pre-intervention time to talk in advance with student | 1 | 0.730 | 0.396 | 0.01 |
| Residual | 51 | | | |

**Estimated marginal means**

| Condition | Adjusted Mean | SE | 95% CI |
|---|---|---|---|
| *Self-Guided* | 4.11 | 0.23 | [3.63, 4.58] |
| *Self-Guided + Peer Discussion* | 3.58 | 0.22 | [3.13, 4.04] |
